# Telomere-to-telomere assembly of the genome of an individual *Oikopleura dioica* from Okinawa using Nanopore-based sequencing

**DOI:** 10.1101/2020.09.11.292656

**Authors:** Aleksandra Bliznina, Aki Masunaga, Michael J. Mansfield, Yongkai Tan, Andrew W. Liu, Charlotte West, Tanmay Rustagi, Hsiao-Chiao Chien, Saurabh Kumar, Julien Pichon, Charles Plessy, Nicholas M. Luscombe

## Abstract

**Background:** The larvacean *Oikopleura dioica* is an abundant tunicate plankton with the smallest (65-70 Mbp) non-parasitic, non-extremophile animal genome identified to date. Currently, there are two genomes available for the Bergen (OdB3) and Osaka (OSKA2016) *O. dioica* laboratory strains. Both assemblies have full genome coverage and high sequence accuracy. However, a chromosome-scale assembly has not yet been achieved.

**Results:** Here, we present a chromosome-scale genome assembly (OKI2018_I69) of the Okinawan *O. dioica* produced using long-read Nanopore and short-read Illumina sequencing data from a single male, combined with Hi-C chromosomal conformation capture data for scaffolding. The OKI2018_I69 assembly has a total length of 64.3 Mbp distributed among 19 scaffolds. 99% of the assembly is in five megabase-scale scaffolds. We found telomeres on both ends of the two largest scaffolds, which represent assemblies of two fully contiguous autosomal chromosomes. Each of the other three large scaffolds have telomeres at one end only and we propose that they correspond to sex chromosomes split into a pseudo-autosomal region and X-specific or Y-specific regions. Indeed, these five scaffolds mostly correspond to equivalent linkage groups of OdB3, suggesting overall agreement in chromosomal organization between the two populations. At a more detailed level, the OKI2018_I69 assembly possesses similar genomic features in gene content and repetitive elements reported for OdB3. The Hi-C map suggests few reciprocal interactions between chromosome arms. At the sequence level, multiple genomic features such as GC content and repetitive elements are distributed differently along the short and long arms of the same chromosome.

**Conclusions:** We show that a hybrid approach of integrating multiple sequencing technologies with chromosome conformation information results in an accurate *de novo* chromosome-scale assembly of *O. dioica*’s highly polymorphic genome. This assembly will be a useful resource for genome-wide comparative studies between *O. dioica* and other species, as well as studies of chromosomal evolution in this lineage.

## Background

Larvaceans (synonym: appendicularians) are among the most abundant and ubiquitous taxonomic groups within animal plankton communities (Alldredge, 1976; Hopcroft and Roff, 1995). They live inside self-built “houses” which are used to trap food particles (Sato *et al.*, 2001). The animals regularly replace houses as filters become damaged or clogged and a proportion of discarded houses with trapped materials eventually sink to the ocean floor. As such larvaceans play a significant role in global vertical carbon flux (Alldredge, 2005).

*Oikopleura dioica* is the best documented species among larvaceans. It possesses several invaluable features as an experimental model organism. It is abundant in coastal waters and can be easily collected from the shore. Multigenerational culturing is possible (Masunaga *et al.*, 2020). It has a short lifecycle of 4 days at 23°C and remains free-swimming throughout its life (Feanux, 1998). As a member of the tunicates, a sister taxonomic group to vertebrates, *O. dioica* offers insights into their evolution (Delsuc *et al.*, 2006).

*O. dioica*’s genome size is 65-70 Mb (Seo *et al.*, 2001; Denoeud *et al.*, 2010), which is one of the smallest among all sequenced animals. Interestingly, genome sequencing of other larvacean species uncovered larger genome sizes, which correlated with the expansion of repeat families (Naville *et al.*, 2019). *O. dioica* is distinguished from other tunicates as it is the only reported dioecious species (Fredriksson and Olsson, 1991) and its sex determination system uses an X/Y pair of chromosomes (Denoeud *et al.*, 2010). The first published genome assembly of *O. dioica* (OdB3, B stands for Bergen) was performed with Sanger sequencing which allowed for high sequence accuracy but limited coverage (Denoeud *et al.*, 2010). The OdB3 assembly was scaffolded with a physical map produced from BAC end sequences, which revealed two autosomal linkage groups and a sex chromosome with a long pseudo-autosomal region (PAR; Denoeud *et al.*, 2010). Recently, a genome assembly for a mainland Japanese population of *O. dioica* (OSKA2016, OSKA denotes Osaka) was published, which displayed a high level of coding sequence divergence compared with the OdB3 reference (Wang *et al.*, 2015; Wang *et al.*, 2020). Although OSKA2016 was sequenced with single-molecule long reads produced with the PacBio RSII technology, it does not have chromosomal resolution.

Historical attempts at karyotyping *O. dioica* by traditional histochemical stains arrived at different chromosome counts, ranging between *n* = 3 (Körner, 1952) and *n* = 8 (Colombera and Fernaux, 1973). In preparation for this study, we karyotyped the Okinawan *O. dioica* by staining centromeres with antibodies targeting phosphorylated histone H3 serine 28 (Liu *et al.*, 2020), and concluded a count of *n* = 3. This is also in agreement with the physical map of OdB3 (Denoeud *et al.*, 2010).

Currently, the method of choice for producing chromosome-scale sequences is to assemble contigs using long reads (~10 kb or more) produced by either the Oxford Nanopore or PacBio platforms, and to scaffold them using Hi-C contact maps (Lieberman-Aiden *et al.*, 2009; Dudchenko *et al.*, 2017). To date, there have been no studies of chromosome contacts in *Oikopleura* or any other larvaceans.

Here, we present a chromosome-length assembly of the Okinawan *O. dioica* genome sequence generated with datasets stemming from multiple genomic technologies and data types, namely long-read sequencing data from Oxford Nanopore, short-read sequences from Illumina and Hi-C chromosomal contact maps (Fig. 1).

**Figure 1:**
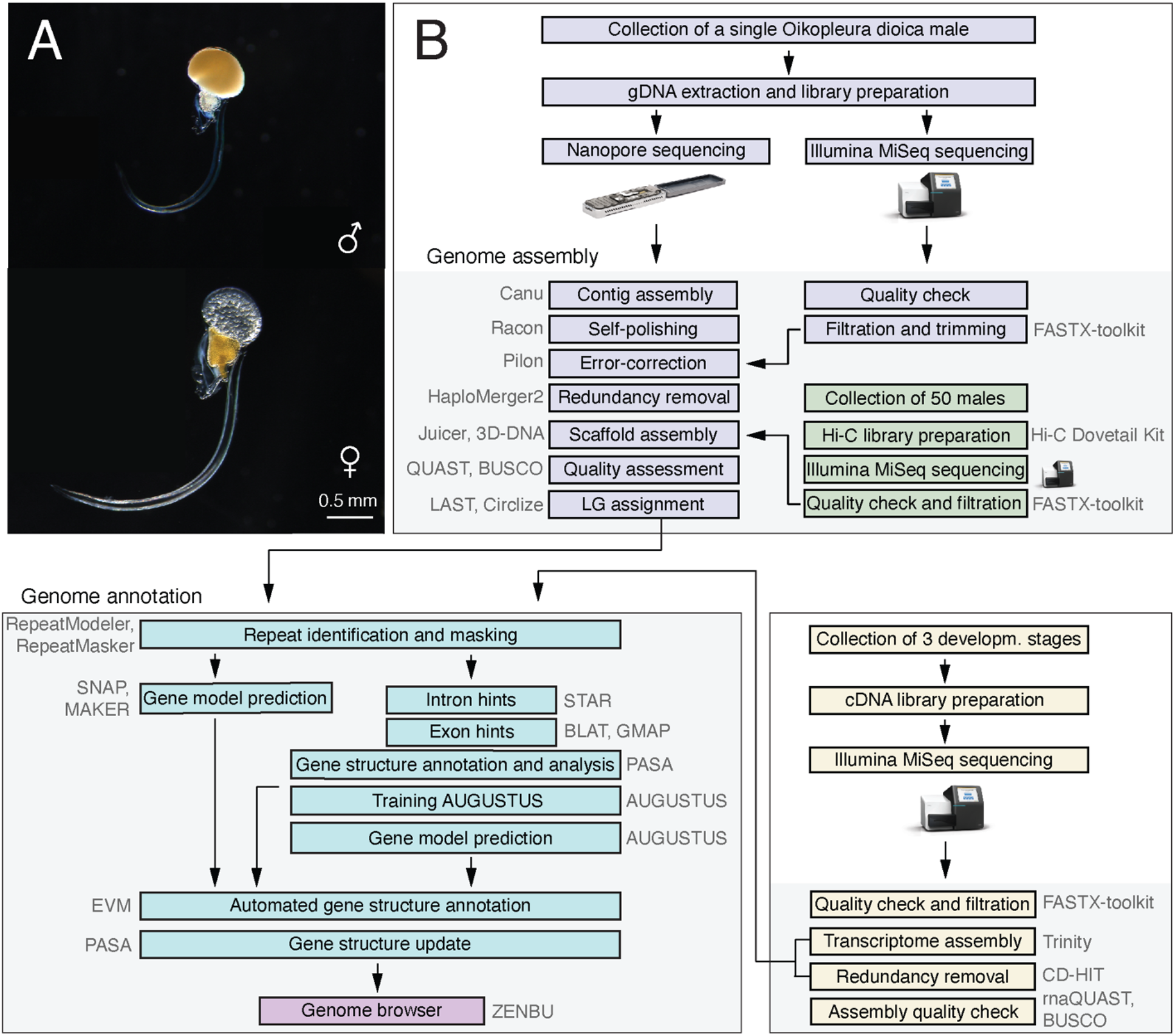
(A) Life images of adult male (top) and female (bottom) *O. dioica*. (B) Genome assembly and annotation workflow that was used to generate the OKI2018_I69 genome assembly.

## Results

### Genome sequencing and assembly

*O. dioica*’s genome is small and highly polymorphic (Denoeud *et al.*, 2010), making assembly of its complete sequence challenging. To reduce the level of variation, we sequenced genomic DNA from a single *O. dioica* male. A low amount of extracted DNA is an issue when working with small-size organisms like *O. dioica.* Therefore, we optimized the extraction and sequencing protocols to allow for low-template input DNA yields of around 200 ng and applied a hybrid sequencing approach using Oxford Nanopore reads to span repeat-rich regions and Illumina reads to correct individual nucleotide errors. The Nanopore run gave 8.2 million reads (221× coverage) with a median length of 840 bp and maximum length of 166 kb (Fig. 2A). Based on k-mer counting of the Illumina reads, the genome was estimated to contain ~50 Mbp (Fig. 2B) – comparable in size to the OdB3 and OSKA2016 assemblies – and a relatively high heterozygosity of ~3.6%. We used the Canu pipeline (Koren *et al.*, 2017) to correct, trim and assemble Nanopore reads, yielding a draft assembly comprising 175 contigs with a weighted median N50 length of 3.2 Mbp. We corrected sequencing errors and local misassemblies of the draft contigs with Nanopore reads using Racon, and then with Illumina reads using Pilon. The initial Okinawa *O. dioica* assembly length was 99.3 Mbp, or ~1.5 times longer than the OdB3 genome at 70.4 Mbp. Merging haplotypes with HaploMerger2 resulted in two sub-assemblies (reference and allelic) of 64.3 Mbp with an N50 of 4.7 Mbp (I69-4). Repeating the procedure on a second individual from the same culture showed overall agreement in assembly lengths, sequences and structures (Fig. 2C).

**Figure 2:**
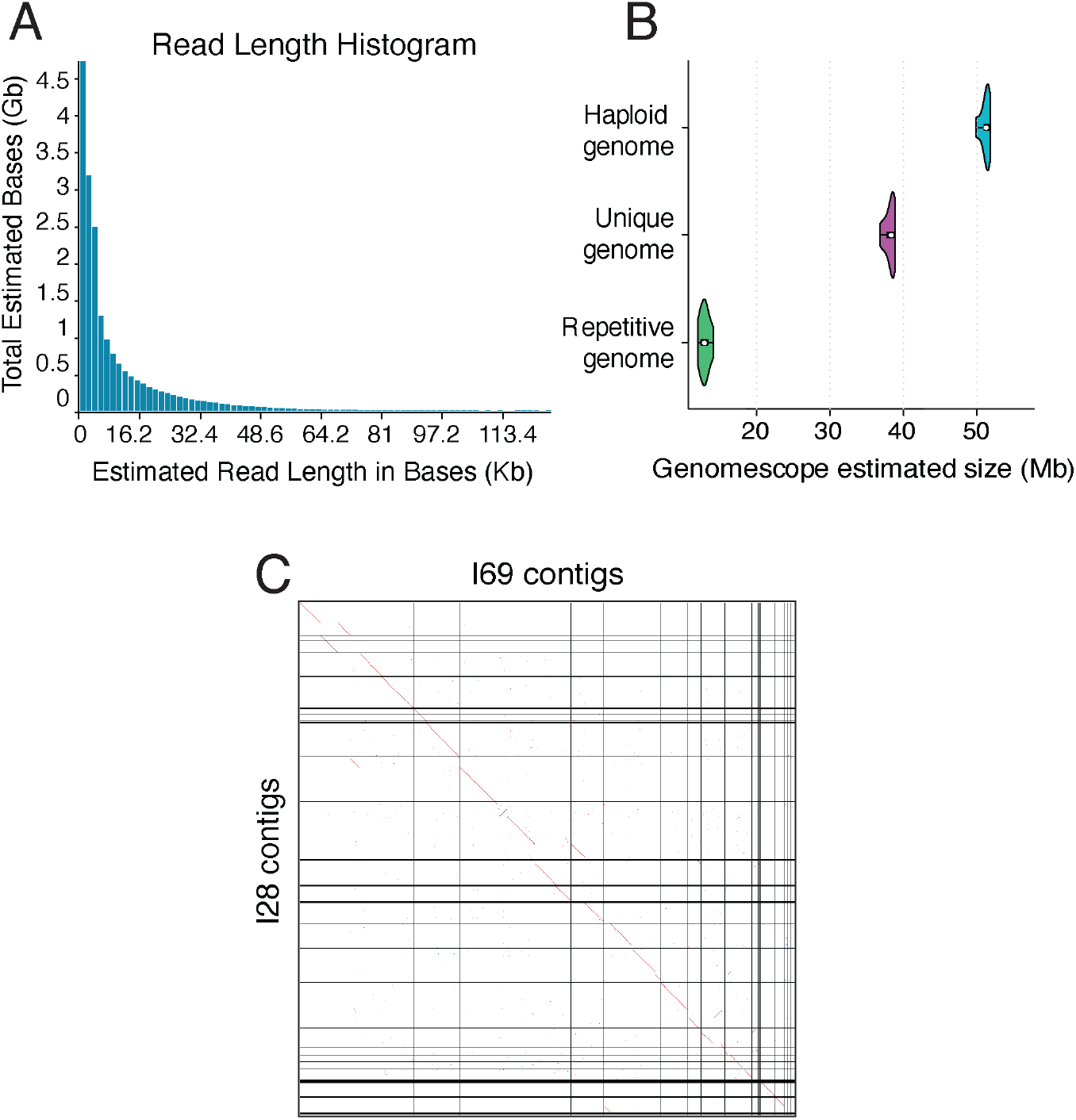
(A) Length distribution of raw Nanopore reads. (B) Estimated total and repetitive genome size based on *k*-mer counting of the Illumina paired-end reads used for assembly polishing. (C) Pairwise genome alignment of the contig assemblies of I69 and I28 *O. dioica* individuals.

To scaffold the genome, we sequenced Hi-C libraries from a pool of ~50 individuals from the same culture. More than 99% of the Hi-C reads could be mapped to the contig assembly. After removing duplicates, Hi-C contacts were passed to the 3D-DNA pipeline to correct major misassemblies, as well as order and orient the contigs. The resulting assembly named consisted of 8 megabase-scale scaffolds containing 99% of the total sequence (Fig. 3A), and 14 smaller scaffolds that account for the remaining 663 Kbp (lengths ranging from 2.9 to 131.6 Kbp). One of the small scaffolds is a draft assembly of mitochondrial genome that we discuss below. Most of the other smaller scaffolds are highly repetitive and might represent unplaced fragments of centromeric or telomeric regions. We annotated telomeres by searching for the TTAGGG repeat sequence and found that most of the megabase-scale scaffolds have single telomeric regions: therefore, we reasoned that they represent chromosome arms. Indeed, pairwise genome alignment to OdB3 identified two syntenic scaffolds for each autosomal linkage group, two for the pseudo-autosomal region (PAR) and one for each sex-specific region. Since we had previously inferred a karyotype of *n* = 3 by immunohistochemistry (Liu *et al.*, 2020), we completed the assembly by pairing the megabase-scale scaffolds into chromosome arms based on their synteny with the OdB3 physical map (Fig. 3B). The final assembly named OKI2018_I69 (Table 1; Suppl. Table 1) comprises telomere-to-telomere assemblies of the autosomal chromosomes 1 (chr 1) and 2 (chr 2). The sex chromosomes are split in pseudo-autosomal region (PAR) and X-specific region (XSR) or Y-specific region (YSR; Fig. 3). We assume that the sex-specific regions belong to the long arm of the PAR, as the long arm does not comprise any telomeric repeats (Fig. 4A).

**Table 1:**
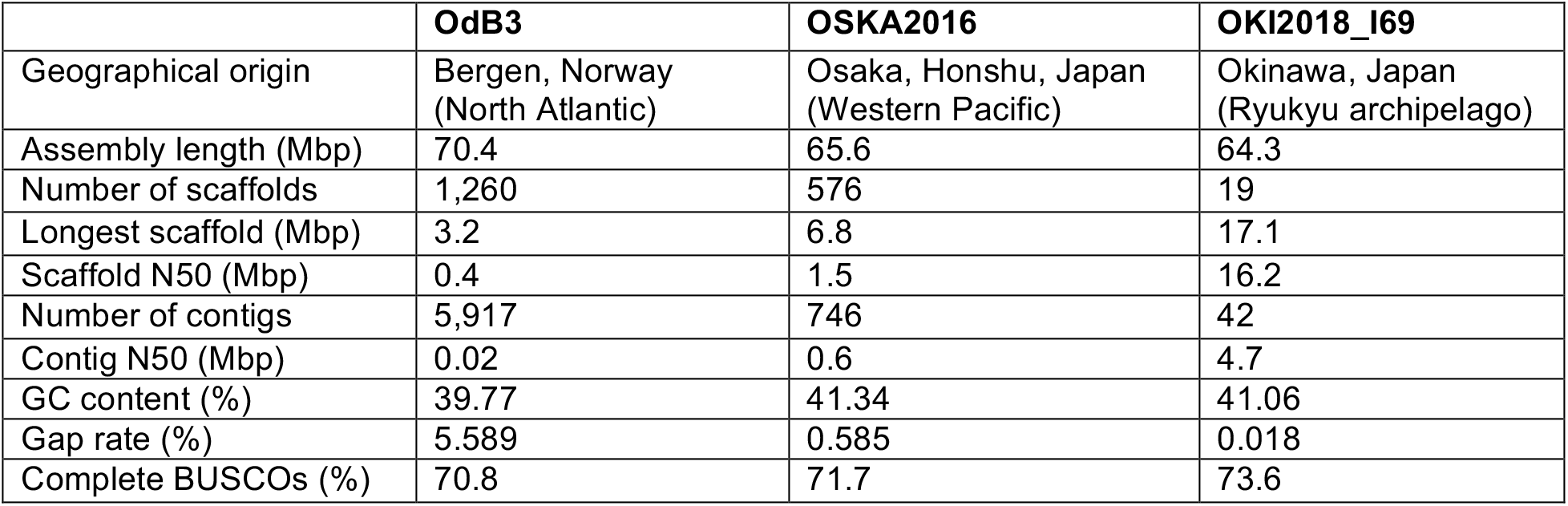
Comparison of the OKI2018_I69 assembly with the previously published *O. dioca* genomes.

**Figure 3:**
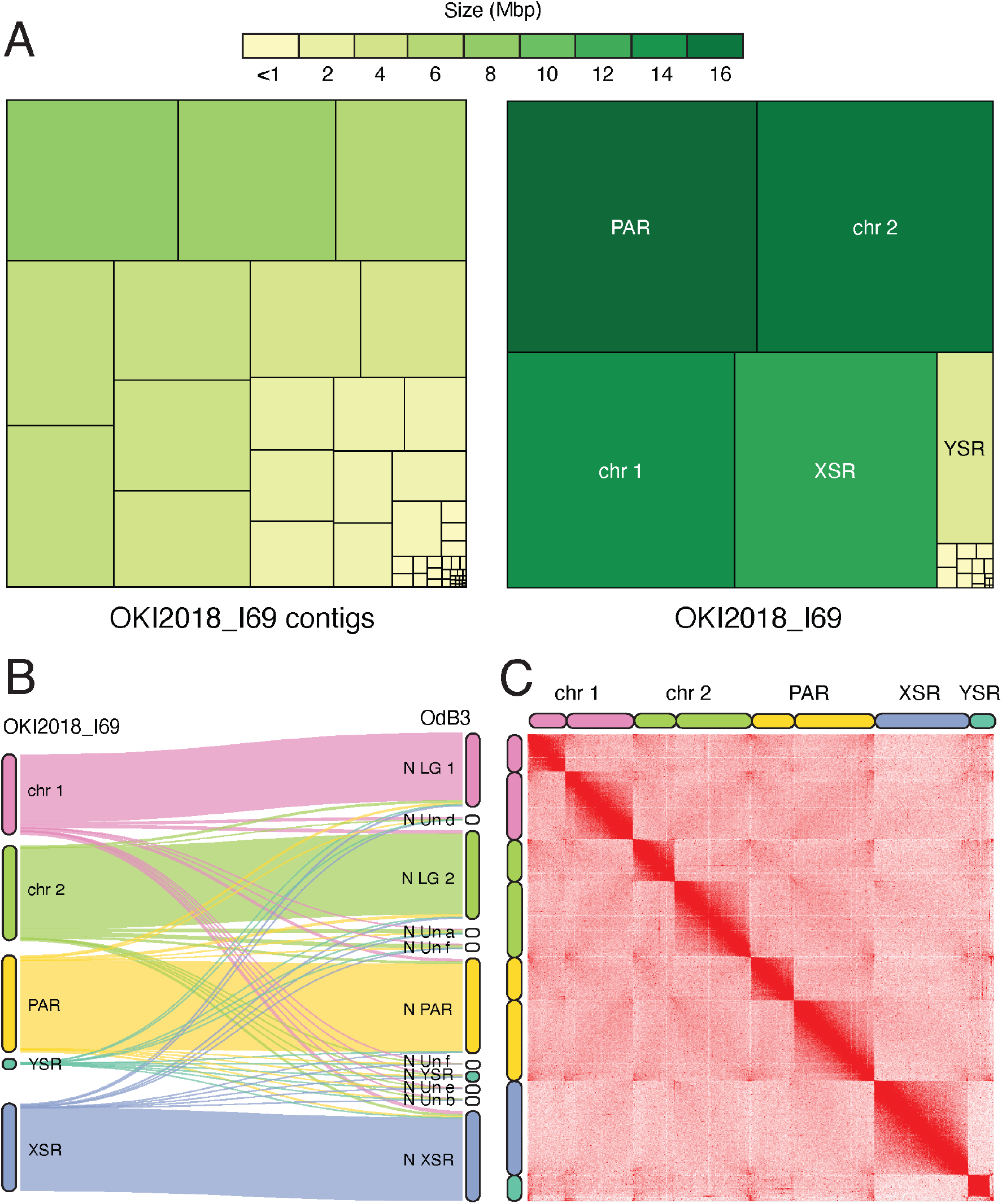
(A) Treemap comparison between the contig (left) and scaffold (right) assemblies of the *O. dioica* genome. Each rectangle represents a contig or a scaffold in the assembly with the area proportional to its length. (B) Comparison between the OKI2018_I69 (left) and OdB3 (right) linkage groups. The Sankey plot shows what proportion of each chromosome in the OKI2018_I69 genome is aligned to the OdB3 linkage groups. (C) Contact matrix generated by aligning Hi-C data set to the OKI2018_I69 assembly with Juicer and 3D-DNA pipelines. Pixel intensity in the contact matrices indicates how often a pair of loci collocate in the nucleus.

**Figure 4:**
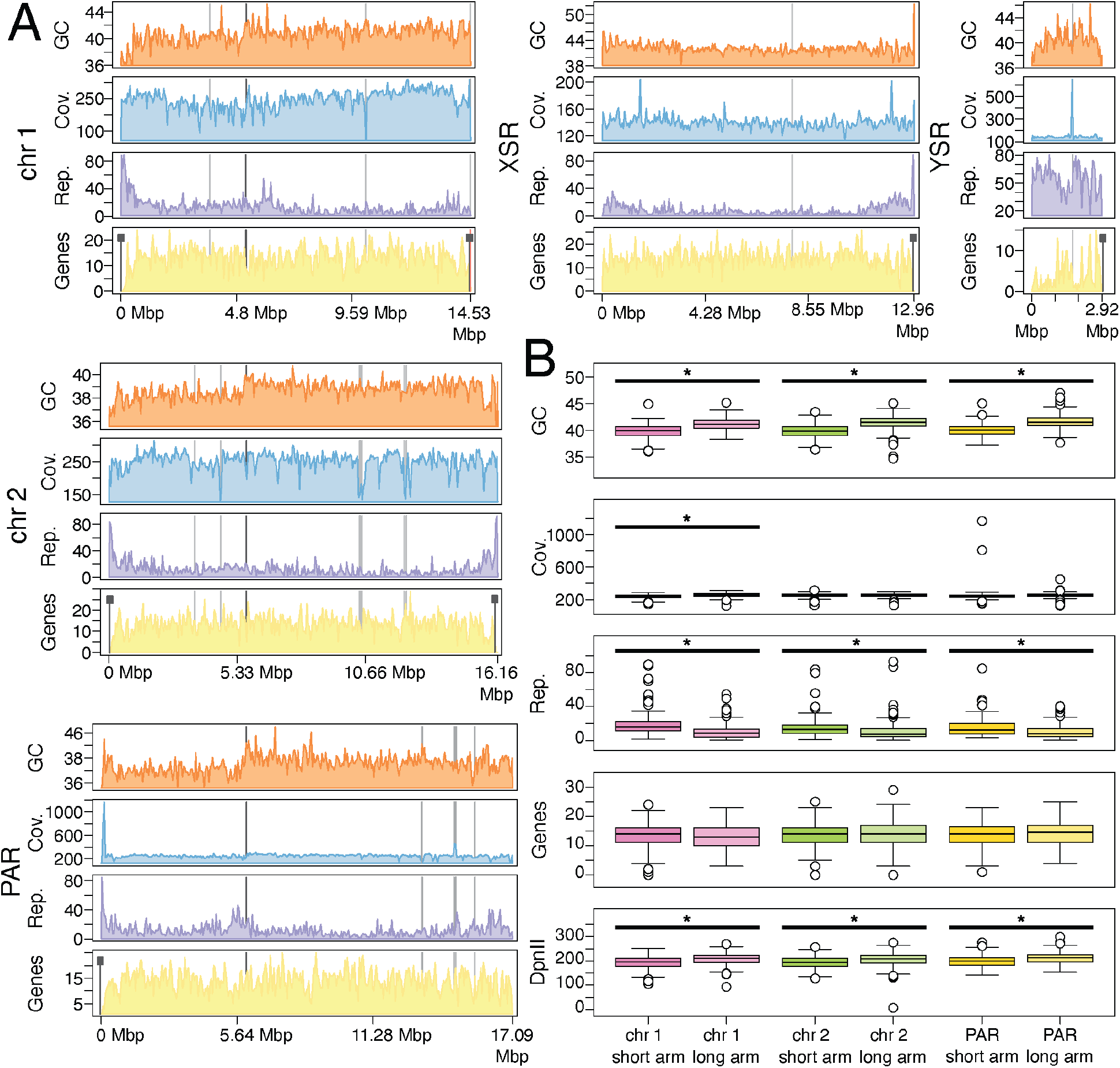
(A) Visualization of sequence properties across chromosomes in the OKI2018_I69 assembly. For each chromosome, 50 Kbp windows of GC (orange), Nanopore sequence coverage (blue), the percent of nucleotides masked by RepeatMasker (purple), and the number of genes (yellow) are indicated. Differences in these sequence properties occur near predicted sites of centromeres and telomeres, as well as between the short and long arms of each non-sex-specific chromosome. Telomeres and gaps in the assembly are indicated with black and grey rectangles, respectively. B) Long and short chromosome arms exhibit significant differences sequence properties, including GC content, repetitive sequence content, and the number of restriction sites recognized by the DpnII enzyme used to generate the Hi-C library.

The genome-wide contact matrix from the Hi-C data (Fig. 3C) shows bright, off-diagonal spots that suggest spatial clustering of the telomeres and centromeres both within the same and across different chromosomes (Dudchenko *et al.*, 2017). The three centromeric regions are outside the sex-specific regions, dividing the PAR and both autosomes into long and short arms. The two sex-specific regions have lower apparent contact frequencies compared with the rest of the assembly which is consistent with their haploid status in males. The chromosome arms themselves show few interactions between each other, even when they are part of the same chromosome.

### Chromosome-level features

The genome contains between 1.4 and 2.6 Mbp of tandem repeats (detected using the tantan and ULTRA algorithms respectively with maximum period lengths of 100 and 2,000). Subtelomeric regions tend to contain retrotransposons or tandem repeats with longer periods. We also found telomeric repeats in smaller scaffolds. A possible explanation is that subtelomeric regions display high heterozygosity, leading to duplicated regions that fail to assemble with the chromosomes. Alternatively, these scaffolds could be peri-centromeric regions containing interstitial telomeric sequences. In some species, high-copy tandem repeats can be utilized to discover the position of centromeric regions (Melters *et al.*, 2013). However, we could not find such regions. Additional experimental techniques such as chromatin immunoprecipitation and sequencing with centromeric markers might be necessary to resolve the centromeres precisely. Therefore, the current assembly skips over centromeric regions, represented as gaps of an arbitrary size of 500 bp in the chromosomal scaffolds.

We studied genome-scale features by visualizing them along whole chromosomes, from the short to long arm, centered on their centromeric regions. Most strikingly, there is a clear difference in sequence content between chromosome arms (Fig. 4; Supp. Table 3). For each chromosome, small arms consistently display depleted GC content and elevated repetitive content compared with the same chromosome’s long arm. Although GC content tends to be weakly negatively correlated with repeat content, it is difficult to ascertain whether repetitive elements tend to drive changes in GC content or if changes in GC content tend to drive the accumulation of repetitive sequence content. In either case, the mechanism behind the marked difference in sequence content between short and large chromosome arms remains unknown. It should be noted that the GC difference between short and long arms also has an effect on the availability of DpnII restriction enzyme recognition sites used for Hi-C library preparation, which recognizes the GC-rich motif/GATC, although this bias is likely insufficient to explain the low degree of intra-chromosomal interaction observed in the Hi-C contact maps.

### Quality assessment using BUSCO

To assess the completeness of our assembly, we searched for 978 metazoan Benchmarking Universal Single-Copy Orthologs (BUSCOs) provided with the BUSCO tool (Simão *et al.*, 2015; Waterhouse *et al.*, 2017; Zdobnov *et al.*, 2017). To increase sensitivity, we trained BUSCO’s gene prediction tool, AUGUSTUS (Hoff and Stanke, 2019), with transcript models generated from RNA-Seq data collected from the same laboratory culture (see below). We detected 73.0% of BUSCOs (Table 1), which is similar to OdB3 and OSKA2016 (Fig. 5A; Suppl. Table 4). All detected BUSCOs except one reside on the chromosomal scaffolds. As the reported fraction of detected genes is lower than for other tunicates such as *Ciona intestinalis* HT (94.6%; Satou *et al.*, 2019) or *Botrylloides leachii* (89%; Blanchoud *et al.*, 2018), we searched for BUSCO genes in the transcriptomic training data (83.0% present) and confirmed the presence of all but one by aligning the transcript sequence to the genome. We then inspected the list of BUSCO genes that were found neither in the genome nor in the transcriptome. Bibliographic analysis confirmed that BUSCO genes related to the peroxisome were lost (Žárský and Tachezy, 2015; Kienle *et al.*, 2016). There are two possible explanations for the remaining missing genes: first is that protein sequence divergence (Berná *et al.*, 2012) or length reduction (Berná and Alvarez-Valin, 2015) in *Oikopleura* complicate detection by BUSCO, and second is gene loss. In line with the possibility of gene loss, most BUSCO genes missing from our assembly are also undetectable in OdB3 and OSKA2016 (Fig. B; Suppl. Table 5). To summarize, the Okinawa assembly achieved comparable detection of universal single-copy conserved orthologs in comparison to previous *O. dioica* assemblies, and consistently undetectable genes may be in fact missing or altered in *Oikopleura*.

**Figure 5:**
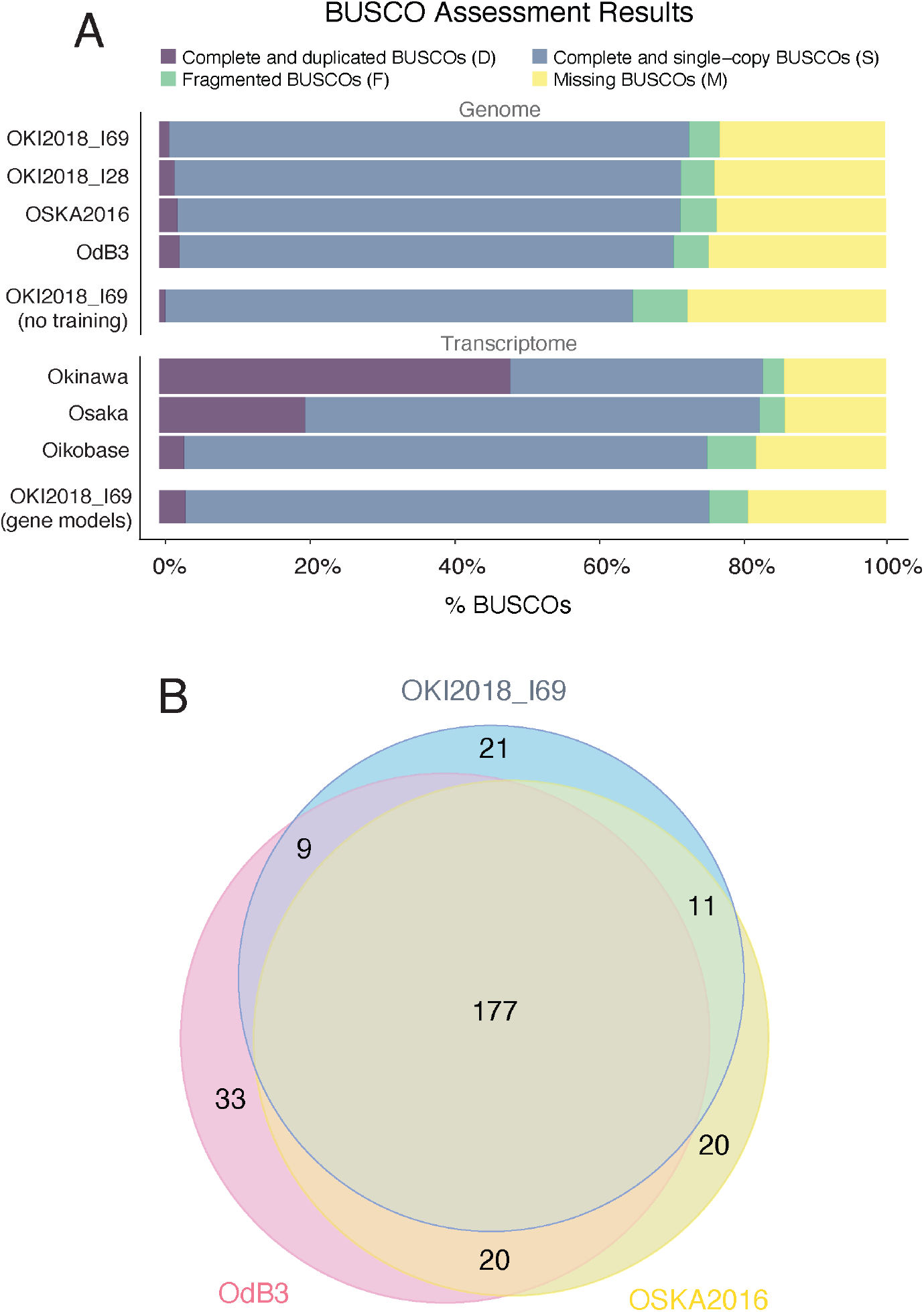
(A) Proportion of BUSCO genes detected or missed in *Oikopleura* genomes and transcriptomes. The search on the OKI2018_I69 assembly was repeated with default parameters (“no training”) to display the effect of AUGUSTUS training. (B) Number of BUSCO genes missing in one or multiple reference genomes.

### Repeat annotation

In order to identify repetitive elements in the OKI2018_I69 genome, we combined the results of several *de novo* repeat detection algorithms and used this custom library as an input to RepeatMasker to identify repeat sequences. Interspersed repeats make up 14.39% of the assembly (9.25 Mbp; Fig. 6), comparable to the 15% reported for OdB3 (Denoeud *et al.*, 2010). Of the annotated elements, the most abundant type is the long terminal repeats (LTRs; ~4.6%) with Ty3/gypsy *Oikopleura* transposons (TORs) dominating 2.97 Mbp of the sequence. Short interspersed nuclear elements (SINEs) make up a smaller portion of the OKI2018_I69 sequence (<0.1%) compared with the OdB3 (0.62%). It has been suggested that SINEs contribute significantly to genome size variation in other oikopleurids (Naville *et al.*, 2019), but further analysis is required to determine whether that is the case at shorter evolutionary distances. Non-LTR LINE/Odin and Penelope-like elements are large components of most oikopleurid genomes (Naville *et al.*, 2019), but they are almost absent in the OKI2018_I69 assembly. Indeed, 44% of the Okinawa *O. dioica* predicted repeats could not be classified through searches against repeat databases and may either represent highly divergent relatives of known repeat classes, or else potentially represent novel repeats specific to Okinawan *O. dioica*.

**Figure 6:**
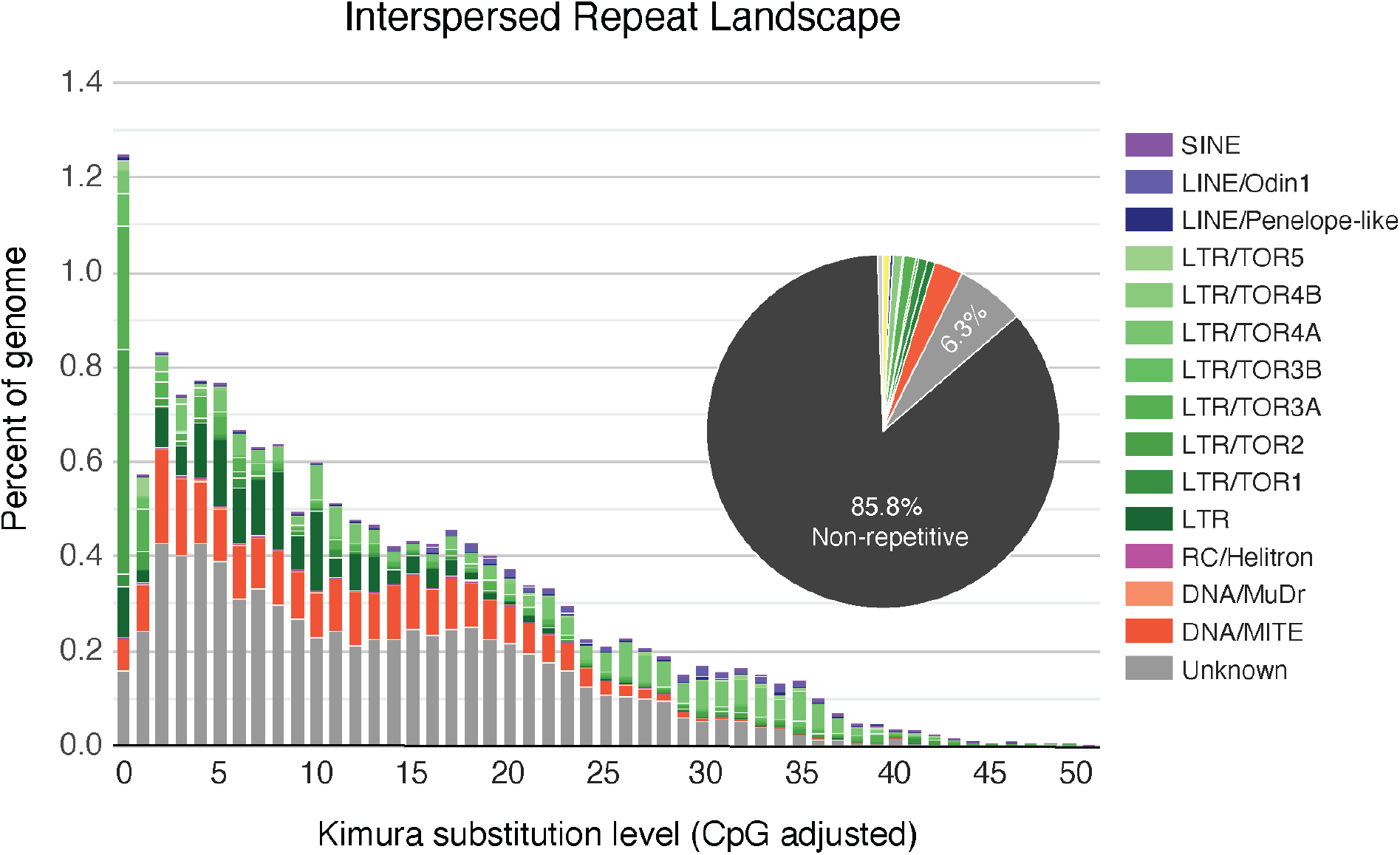
Analysis of repetitive elements. The repeat landscape and proportions of various repeat classes in the genome are indicated and color-coded according to the classes shown on the right side of the figure. The non-repetitive fraction of the genome is shown in black.

### Gene annotation

We annotated the OKI2018_I69 assembly using *ab initio* and RNA-Seq-based gene predictions. The different predictions were refined and merged with EVidenceModeler using the Okinawan transcriptome and Bergen ESTs and proteins as additional support. To predict alternative isoforms and update the models, we ran the PASA annotation pipeline that yielded 18,485 transcript isoforms distributed among 16,936 protein-coding genes. The number of predicted genes for the OKI2018_I69 is lower than what was reported for OdB3 (18,020; Denoeud *et al.*, 2010) and OSKA2016 (18,743; Wang *et al.*, 2020). The rest of the genes are either lost from the Okinawan *O. dioica* genome or were not assembled and/or annotated with our pipeline. On the other side, higher number of genes might be artifact of the OdB3 and OSKA2016 annotations. The completeness of our annotation compares to the genome: BUSCO recovered 75.7% complete and 5.3% fragmented metazoan genes (Fig. 5A). Like in the OdB3 assembly, gene density is very high at one gene per 3.69 Kbp. Genes have very short introns (median length at 46 bp) and intergenic spaces (average length 1,206 bp) (Table 2). Therefore, overall genomic features seem to be conserved among *O. dioica* population despite large geographic distance.

**Table 2:**
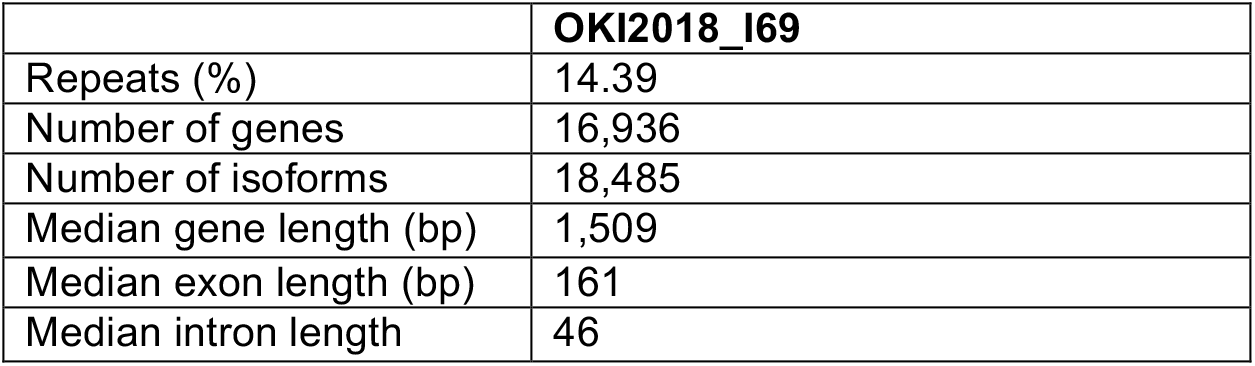
Overall characterization of the OKI2018_I69 genome assembly

The ribosomal DNA gene encoding the precursor of the 18S, 5.8S and 28S rRNAs occurs as long tandem repeats that form specific chromatin domains in the nucleolus. We identified 4 full tandem copies of the rDNA gene at the tip of the PAR’s short arm, separated by 8,738 bp (median distance). As this region has excess coverage of raw reads, and since assemblies of tandem repeats are limited by the read length (99% of Nanopore reads in our data are shorter than 42,842 bp), we estimate that the real number of the tandem rDNA copies could range between 5 (MiSeq) and 25 times (Nanopore) larger. Between or flanking the rDNA genes, we also found short tandem repeats made of 2 to 3 copies of a 96-bp sequence. This tandem repeat is unique to the rDNA genes and to our reference and draft genomes, and was not found in the OdB3 reference nor in other larvacean genomes. The 5S rRNA is transcribed from loci distinct to the rDNA gene tandem arrays. In *Oikopleura*, it has the particularity of being frequently associated with the spliced leader (SL) gene and to form inverted repeats present in more than 40 copies (Ganot *et al.*, 2004). We found 27 copies of these genes on every chromosomal scaffold except YSR, 22 of which were arranged in inverted tandem repeats. Altogether, we found in our reference genome one rDNA gene repeat region assembled at the end of a chromosome short arm. This sequence might provide useful markers for phylogenetic studies in the future.

### Draft mitochondrial genome scaffold

We identified a draft mitochondrial genome among the smaller scaffolds, chrUn_12, by searching for mitochondrial sequences using the Cox1 protein sequence and the ascidian mitochondrial genetic code (Pichon *et al.*, 2019). Automated annotation of this scaffold using the MITOS2 server detected the coding genes *cob*, *cox1*, *nad1*, *cox3*, *nad4*, *cox2*, and *atp6* (Fig. 7A), which are the same as in Denoeud *et al.*, 2010 except for the *nd5* gene that is missing from our assembly. The open reading frames are often interrupted by T-rich regions, in line with Denoeud *et al.* (2010). However, we cannot rule out the possibility that these regions represent sequencing errors, as homopolymers are difficult to resolve with the Nanopore technology available in 2019. The *cob* gene is interrupted by a long non-coding region, but this might be a missassembly. Indeed, an independent assembly using the `flye` software (Kolmogorov *et al.*, 2019) with the `--meta` option to account for differential coverage also produced a draft mitochondrial genome, but its non-coding region was ~2 kbp longer. Moreover, a wordmatch dotplot shows tandem repeats in this region (Fig. 7B), and thus this region is prone to assembly errors, especially with respect to the number of repeats. Altogether, the draft contig produced in our assembly shows as a proof of principle that sequencing reads covering the mitochondrial genome alongside the nuclear genome can be produced from a single individual, although it may need supporting data such as targeted resequencing in order to be properly assembled.

**Figure 7:**
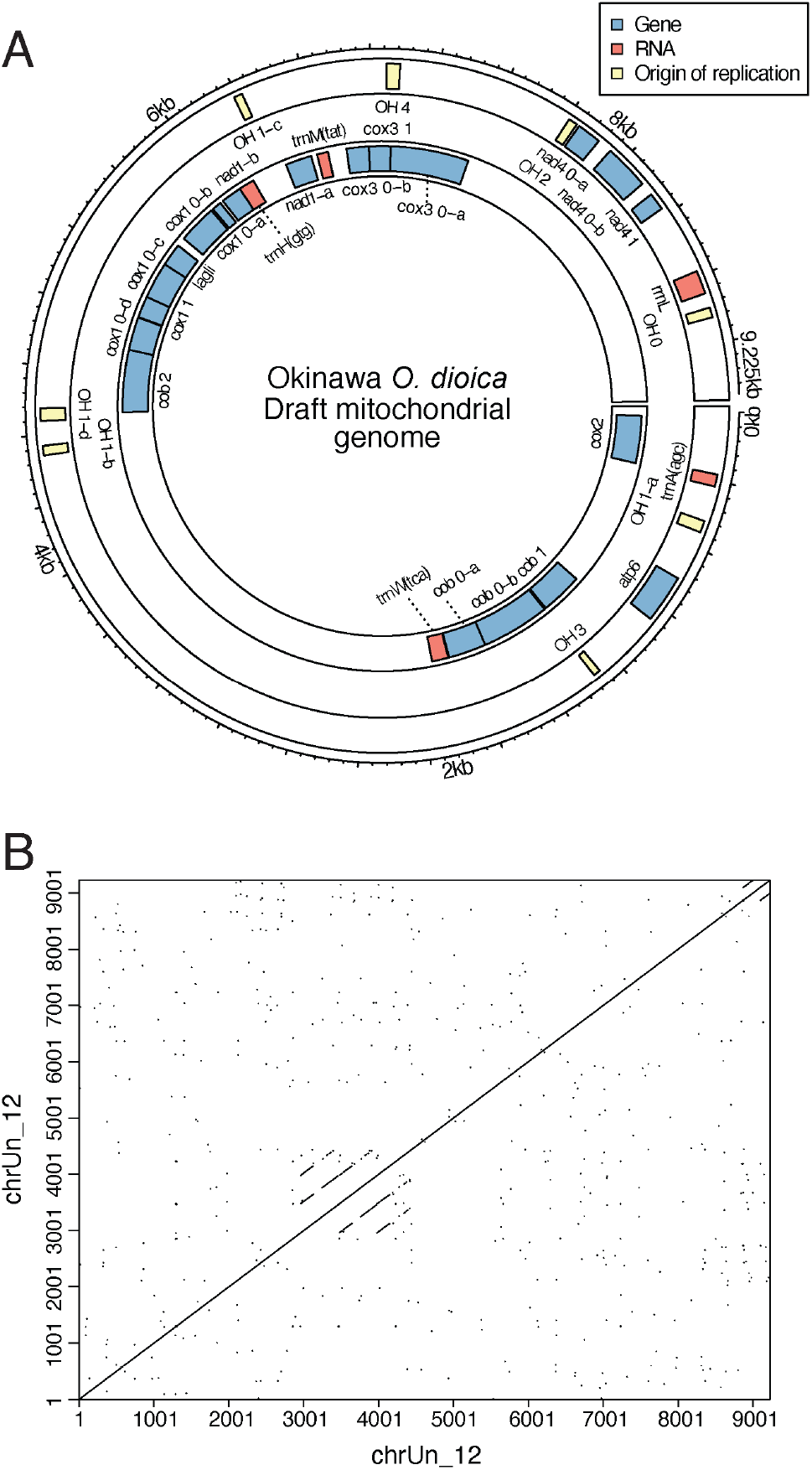
(A) Predicted gene annotation of the draft mitochondrial genome sequence. (B) Self-similarity plot of the draft mitochondrial genome sequence. A tandem repeat can be seen, which complicates the complete assembly of the mitochondrial genome from whole-genome sequencing data.

## Discussion

### OKI2018_I69 assembly quality

Previously, different techniques have been used to sequence and assemble *O. dioica* genomes which have produced assemblies of varying quality. The Sanger-based OdB3 sequence was published in 2010 (Denoeud *et al.*, 2010). Due to limitations in sequencing technologies at the time, it is highly fragmented, comprising 1,260 scaffolds with an N50 of 0.4 Mbp. The recently released OSKA2016 assembly was generated from long-read PacBio data and, therefore, has a larger N50 and fewer scaffolds (Table 1, Wang *et al.* 2020). Both assemblies have high sequence quality and nearly full genome coverage, but neither of them contains resolved chromosomes. However, Denoeud *et al.* (2010) released a physical map calculated for OdB3 from BAC end sequences that comprises five linkage groups (LGs): two autosomal LGs, one pseudo-autosomal region of sex chromosomes, and two sex specific regions (X and Y).

The use of reference chromosome information from a closely related species to order contigs or scaffolds into chromosome-length sequences is a common way to generate final genome assemblies (*Drosophila* 12 Genomes Consortium, 2007). However, this approach precludes discovery of structural variances. In our study, we first assembled long Nanopore reads *de novo* into contigs that we ordered and joined into megabase-scale scaffolds using long-range Hi-C data. The synteny-based approach with OdB3’s linkage groups as a reference was only required to guide final pairing of chromosome arms into single scaffolds of chr 1, chr 2 and PAR, as we found that these scaffolds mostly align to one of the autosomal LGs or PAR. Therefore, any potential assembly errors in OdB3 would not be transferred to our assembly. Apart from these syntenic relationships, our karyotyping results and the count of three centromeres on the Hi-C contact map supports the presence of three pairs of chromosomes in the Okinawan *O. dioica*. However, there is a possibility that chromosome arms might have been exchanged between chromosomes in the Okinawan population. Additional experimental evidence is needed to confirm the pairing of chromosome arms, such as data generated by the Omni-C method which does not rely on restriction enzyme fragmentation.

Our synteny-based scaffolding assumes that animals collected from the Atlantic and Pacific oceans are from the same species and conserve these chromosomal properties. However, there are visible differences in gene number and repeat content compared with the OdB3 and OSKA2016. *O. dioica* is distributed all over the world, and all the populations are classified as a single species owing to the lack of obvious morphological differences and limited understanding of population structure. However, the short life span of *O. dioica* combined with limited mobility and high mutation rate contribute to an accelerated genome evolution that might have led to multiple speciation events. Sequence polymorphism was previously noted when comparing the OdB3 genome to genomic libraries of a laboratory strain collected on the North American Pacific coast (Denoeud *et al.*, 2010), and more recently when comparing OdB3 to OSKA2010 (Wang *et al.*, 2015; Wang *et al.*, 2020). Further work will be needed to elucidate the relation of the Okinawan population to the North Atlantic and North Pacific ones.

### Inter-arm contacts

The sequence of *O. dioica*’s chromosomes and their contact map suggest that chromosome arms may be the fundamental unit of synteny in larvaceans. Hi-C contact matrices in vertebrates typically display greater intra-chromosomal than inter-chromosomal interactions. A similar pattern was reported in the tunicate *Ciona robusta* (also known as *intestinalis* type A; Satou *et al.*, 2019) and the lancelet *Branchiostoma floridae* (Simakov, Marlétaz, *et al.*, 2020). By comparison, in flies and mosquitoes, the degree of contacts between two arms of the same chromosome appear to be reduced but nonetheless more frequent than between different chromosomes (Dudchenko *et al.*, 2017). Indeed, in *Drosophila*, the chromosome arms – which are termed Muller elements owing to studies with classical genetics (Schaeffer, 2018) – are frequently exchanged between chromosomes across speciation events. *O. dioica*’s genome shares with fruit flies its small size and small number of chromosomes. However, small chromosome size is also seen in the tunicate *Ciona robusta*, which has 14 meta- or sub-meta-centric pairs (Shoguchi *et al.*, 2005), with an average length of ~8 Mbp (Satou *et al.*, 2019) that exhibit a more extensive degree of contacts, particularly for intra-chromosomal interactions across the centromeres (Satou *et al.*, 2019). As we prepared our Hi-C libraries from adult animals, where polyploidy is high (Ganot and Thompson, 2002), we cannot rule out that it could be a possible cause of the low inter-arm interactions in our contact matrix. Further studies such as investigations of other developmental stages will be needed to elucidate the mechanism at work for the similarity between *O. dioica* and insect’s chromosome contact maps.

### Visualization and access

We prepared a public view of our reference genome in the ZENBU browser (Severin *et al.*, 2014), displaying tracks for our gene models, *in silico*-predicted features such as repeats and non-coding RNAs, or syntenies with other *Oikopleura* genomes. To facilitate the study of known genes, we screened the literature for published sequences (Suppl. Table 6) and mapped them to the genome with a translated alignment. The ZENBU track for these alignments is searchable by gene name, accession number and PubMed identifier. Chromosome-level visualization of this track shows that the genes studied so far are distributed evenly on each chromosome, except for the repeat-rich YSR (Figure 8). In line with the observed loss of synteny in the Hox genes noted in *Oikopleura* (Seo *et al.*, 2004), we did not see apparent clustering of genes by function or relatedness. The view of the OKI2018_I69 genome assembly can be found here: https://fantom.gsc.riken.jp/zenbu/gLyphs/#config=nPfav_juDdOmIG2t4LkJ1D;loc=OKI2018_I69_1.0::chr1:1..100000+.

**Figure 8:**
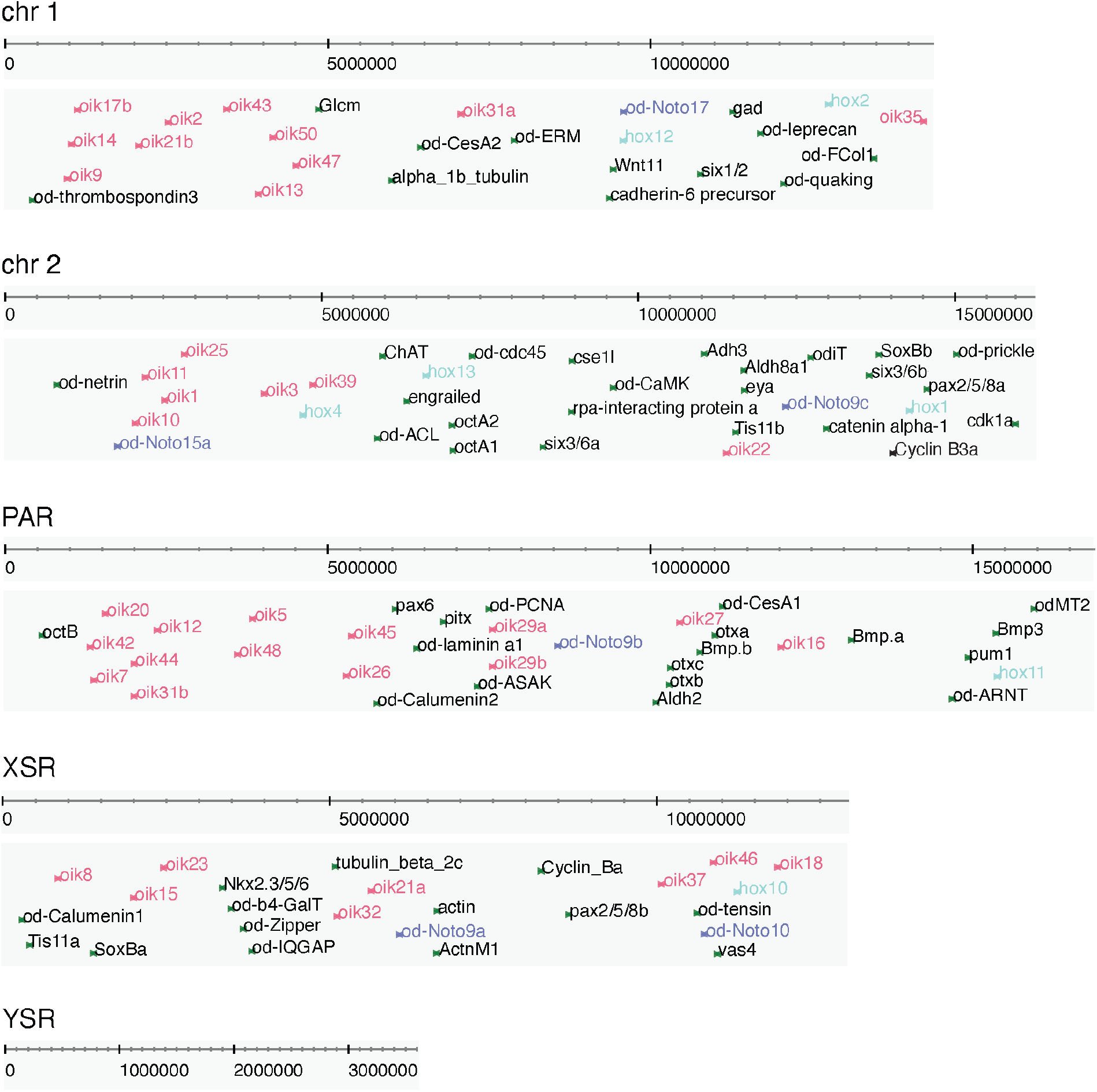
Genomic locations of various oikopleurid gene homologs searchable by name and PubMed identifiers in the ZENBU genome browser. Colours indicate genes from the same family.

## Conclusions

We demonstrated that a combination of long- and short-read sequencing data from a single animal, together with the long-range Hi-C data and the use of various bioinformatic approaches can result in a high-quality *de novo* chromosome-scale assembly of *O. dioica*’s highly polymorphic genome. However, further work is needed to properly resolve the polymorphisms into separated haplotypes using a different approach, such as trio-binning. We believe that the current version of the assembly will serve as an essential resource for a broad range of biological studies, including genome-wide comparative studies of *Oikopleura* and other species, and provides insights into chromosomal evolution.

## Methods

### *Oikopleura* sample and culture

Wild live specimens were collected from Ishikawa Harbor (26°25’39.3”N 127°49’56.6”E) by a hand-held plankton net and returned to the lab for culturing (Masunaga *et al.*, 2020). A typical generation time from hatchling to fully mature adult is 4 days at 23°C for the Okinawan *O. dioica*. Individuals I28 and I69 were collected at generation 44 and 47, respectively.

### Isolation and sequencing of DNA

Staged fully mature males were collected prior to spawning. These were each washed with 5 ml filtered autoclaved seawater (FASW) for 10 min three times before resuspension in 50 μl 4 M guanidium isothiocyanate, 0.5% SDS, 50 mM sodium citrate and 0.05% v/v 2-mercaptoethanol. This was left on ice for 30 min before being precipitated with 2 volumes of ice-cold ethanol and centrifuged at 14,000 rpm 4°C for 20 min. The pellet was washed with 1 ml of 70% cold-ethanol, centrifuged at 14,000 rpm 4°C for 5 min and air dried briefly before resuspension in 200 μl 100 mM NaCl, 25 mM EDTA, 0.5% SDS and 10 μg/ml proteinase K. The lysates were incubated overnight at 50°C. The next morning, the total nucleic acids were first extracted and then back-extracted once more with chloroform:phenol (1:1). Organic and aqueous phases were resolved by centrifugation at 13,000 rpm for 5 min for each extraction; both first and back-extracted aqueous phases were collected and pooled. The pooled aqueous phase was subjected to a final extraction with chloroform and spun down as previously described. The aqueous fraction was then removed and precipitated by centrifugation with two volumes of cold ethanol and 10 μg/ml glycogen; washed with 1 ml of cold 70% ethanol and centrifuged once more as previously described. The resulting pellet was allowed to air-dry for 5 min and finally resuspended in molecular biology grade H_2_O for quantitation using a Qubit 3 Fluorometer (Thermo Fisher Scientific, Q32850), and the integrity of the genomic DNA was validated using Agilent 4200 TapeStation (Agilent, 5067-5365).

Isolated genomic DNA used for long-reads on Nanopore MinION platform were processed with the Ligation Sequencing Kit (Nanopore LSK109) according to manufacturer’s protocol, loading approximately 200 ng total sample per R9.4 flow-cell. Raw signals were converted to sequence files with the Guppy proprietary software (model “template_r9.4.1_450bps_large_flipflop”, version 2.3.5). Approximately 5 ng was set aside for whole genome amplification to perform sequencing on Illumina MiSeq platform, using the TruePrime WGA Kit (Sygnis, 370025) according to manufacturer’s protocol. Magnetic bead purification (Promega, NG2001) was employed for all changes in buffer conditions required for enzymatic reactions and for final buffer suitable for sequencing system. Approximately 1 μg of amplified DNA was sequenced by our core sequencing facility with a 600-cycle MiSeq Reagent Kit v3 (Illumina, MS-102-3003) following the manufacturer’s instructions. These Illumina runs were used for polishing and error checking of Nanopore runs.

### Hi-C library preparation

50 fully matured males were rinsed 3 times for 10 min each by transferring from well to well in a 6-well plate filled with 5 ml FASW. Rinsed animals were combined in a 1.5 ml microcentrifuge tube. Tissues were pelleted for 10 min at 12,000 rpm and leftover FASW was discarded. A Hi-C library was then prepared by following the manufacturer’s protocol (Dovetail, 21004). Briefly, tissues were cross-linked for 20 min by adding 1 ml 1× PBS and 40.5 μl 37% formaldehyde to the pellet. The tubes were kept rotating to avoid tissue settle during incubation. Cross-linked DNA was then blunt-end digested with DpnII (Dovetail) to prepare ends for ligation. After ligation, crosslinks were reversed, DNA was purified by AMPure XP Beads (Beckman, A63880) and quantified by Qubit 3 Fluorometer (Thermo Fisher Scientific, Q10210). The purified DNA was sheared to a size of 250-450 bp by sonication using a Covaris M220 instrument (Covaris, Woburn, MA) with peak power 50 W, duty factor 20, and cycles/burst 200 times for 65 s. DNA end repair, adapter ligation, PCR enrichment, and size selection were carried out by using reagents provided with the kit (Dovetail, 21004). Finally, the library was checked for quality and quantity on an Agilent 4200 TapeStation (Agilent, 5067-5584) and a Qubit 3 Fluorometer. The library was sequenced on a MiSeq (Illumina, SY-410-1003) platform using a 300 cycles V2 sequencing kit (Illumina, MS-102-2002), yielding 20,832,357 read pairs.

### Genome size estimation

Jellyfish (Marçais, Kingsford, 2011) was used to generate k-mer count profiles for various values of *k* (17, 21, 25, 29, 33, 37, and 41) based on the genome-polishing Illumina MiSeq reads, with a maximum k-mer count of 1000. These k-mer profiles were subsequently used to estimate heterozygosity and genome size parameters using the GenomeScope web server (http://qb.cshl.edu/genomescope/).

### Filtering of Illumina MiSeq raw reads

Before using at different steps, all raw Illumina reads were quality-filtered (-q 30, -p 70) and trimmed on both ends with the FASTX-Toolkit v0.0.14 (http://hannonlab.cshl.edu/fastx_toolkit/index.html). The quality of the reads before and after filtering were checked with FASTQC v0.11.5 (Andrews *et al.*, 2010). Read pairs that lacked one of the reads after the filtering were discarded in order to preserve paired-end information.

### Genome assembly

Genome assembly was conducted with the Canu pipeline v1.8 (Koren *et al.*, 2017) and 32.3 Gb (~221.69×) raw Nanopore reads (correctedErrorRate=0.105, minReadLength=1000). The resulting contig assembly was polished three times with Racon v1.2.1 (Vaser *et al.*, 2017) using Canu-filtered Nanopore reads. Nanopore-specific errors were corrected with Pilon v1.22 (Walker *et al.*, 2014) using filtered 150-bp paired-end Illumina reads (~99.7×). Illumina reads were aligned to the Canu contig assembly with BWA v0.7.17 (Li *et al.*, 2013) and the corresponding alignments were provided as input to Pilon. Next, one round of the HaploMerger2 processing pipeline (Huang *et al.*, 2017) was applied to eliminate redundancy in contigs and to merge haplotypes.

Contigs were joined into scaffolds based on long-range Hi-C Dovetail™ data using Juicer v1.6 (Durand *et al.*, 2016) and 3D de novo assembly (3D-DNA; Dudchenko *et al.*, 2017) pipelines. The megabase-scale scaffolds were joined into pairs of chromosome arms based on their synteny with the OdB3 physical map (see below). The candidate assembly was visualized and reviewed with Juicebox Assembly Tools (JBAT) v1.11.08 (https://github.com/aidenlab/Juicebox; Durand and Robinson 2016).

Whole-genome alignment between OKI2018_I69 and OdB3 assemblies was performed using LAST v1066 (Kiełbasa *et al.*, 2011). The sequence of OdB3 linkage groups were reconstructed as defined in the Supplementary Figure 2 in Denoeud et al. 2010. The resulting alignments were post-processed in R with a custom script (https://github.com/oist/oikGenomePaper) and visualized using the R package “networkD3” (“sankeyNetwork” function). The color scheme for chromosomes was adopted from R Package RColourBrewer, “Set2”.

The final assembly was checked for contamination by BLAST searches against the NCBI non-redundant sequence database. 12 smaller scaffolds were found to have strong matches to bacterial DNA (Suppl. Table 2), as well as possessing significantly higher Nanopore sequence coverage (>500×) than the rest of the assembly, and were therefore removed from the final assembly.

The completeness and quality of the assembly were checked with QUAST v5.0.2 (Gurevich *et al.*, 2013) and by searching for the set of 978 highly conserved metazoan genes (OrthoDB version 9.1; Zdobnov *et al.*, 2017) using BUSCO v3.0.2 (Simão *et al.*, 2015; Waterhouse *et al.*, 2017). The --sp option was set to match custom AUGUSTUS parameters (Hoff and Stanke, 2019) trained using the Trinity transcriptome assembly (see below) split 50% / 50% for training and testing.

### Repeat masking and transposable elements

A custom library of repetitive elements (RE) present in the genome assembly was built with RepeatModeler v2.0.1 that uses three -*de novo* -repeat finding programs: RECON v1.08, RepeatScout v1.0.6 and LtrHarvest/Ltr_retriever v2.8. In addition, MITE-Hunter v11-2011 (Han and Wessler, 2010) and SINE_Finder (Wenke and Torsten, *et al.*, 2011) were used to search for MITE and SINE elements, respectively. The three libraries were pooled together as input to RepeatMasker v4.1.0 (Smit, Hubley and Green, 2015) to annotate and soft-mask these repeats in the genomic sequence. Resulting sets of REs were annotated by BLAST searches against RepeatMasker databases and sequences of transposable elements published for different oikopleurids (Naville *et al.*, 2019).

Tandem repeats were detected using two different programs, tantan (Frith, 2011) and ULTRA (Olson and Wheeler, 2018) using two different maximal period lengths (100 and 2000). Version 23 of tantan was used with the parameters -f4 (output repeats) and -w100 or 2000 (maximum period length). ULTRA version 0.99.17 was used with -mu 2 (minimum number of repeats) -p 100 or 2000 (maximum period length) and -mi 5 -md 5 (maximum consecutive insertions or deletions). ULTRA detected more tandem repeats than tantan, but its predictions include more than 90% of tantan’s. Both tools detected *O. dioica*’s telomeric tandem repeat sequence, which is TTAGGG as in other chordates (Schulmeister *et al.*, 2007).

### Developmental staging, isolation and sequencing of mRNA, transcriptome assembly

Mixed stage embryos, immature adults (3 days after hatching) and adults (4 days after hatching) were collected separately from our on-going laboratory culture for RNA-Seq analysis. Eggs were washed three times for 10 min by moving eggs along with micropipette from well to well in a 6-well dish each containing 5 ml of FASW and left in a fresh well of 5 ml FASW in the same dish. These were stored at 17°C and set aside for fertilization. Matured males, engorged with sperm, were also washed 3 times in FASW. Still intact mature males were placed in 100 μl of fresh FASW and allowed to spawn naturally. Staged embryos were initiated by gently mixing 10 μl of the spawned male sperm to the awaiting eggs in FASW at 23°C. Generation 30 developing embryos at 1 h and 3 h post-fertilization were visually verified by dissecting microscope and collected as a pool for the mixed staged embryo time point. Immature adults at generation 31 and sexually differentiated adults at generation 30 were used for the 2 adult staged time points. All individuals for each time point were pooled and washed with FASW three times for 10 min. Total RNA was extracted and isolated with RNeasy Micro Kit (Qiagen, 74004) and quantitated using Qubit 3 Fluorometer (Thermo Fisher Scientific, Q10210). Additional quality control and integrity of isolated total RNA was checked using Agilent 4200 TapeStation (Agilent, 5067-5576). Further processing for mRNA selection was performed with Oligo-d(T)25 Magnetic Beads (NEB, E7490) and the integrity of the RNA was validated once more with Agilent 4200 TapeStation (Agilent, 5067-5579). Adapters for the creation of DNA libraries for the Illumina platform were added per manufacturer’s guidance (NEB, E7805) as were unique indexed oligonucleotides (NEB, E7600) to each of the 3 staged samples. Each cDNA library was sequenced paired-end with a 300-cycle MiSeq Reagent Kit v2 (Illumina, MS-102-2002) loaded at approximately 12 pM.

After quality assessment and data filtering (see Filtering of Illumina MiSeq raw reads), Illumina RNA-Seq reads were pooled together and *de novo* assembled with Trinity v2.8.2 (Grabherr *et al.*, 2011). Redundancy in the transcriptome assembly was removed by CD-HIT v4.8.1 (Li and Godzik, 2006) with a cut-off value of 95% identity. The quality and completeness of the transcriptome assembly was verified with rnaQUAST v1.5.1 (Bushmanova *et al.*, 2016) and BUSCO.

### Gene prediction and annotation

Gene models were predicted using both AUGUSTUS v3.3 (Stanke et al. 2006) and the MAKER pipeline v3.01.03 (Cantarel *et al.*, 2008). AUSGUSTUS was trained following the Hoff and Stanke protocol (2019) with the initial RNA-Seq reads and transcriptome assembly used as intron and exon hints, correspondingly. Transcript models were generated with the PASA pipeline v20140417 (Haas *et al.*, 2003) using BLAT v36 and GMAP v2018-02-12 to align transcripts to the genome. RNA-Seq reads were mapped to the genome with STAR v2.0.6a (Dobin *et al.*, 2013). Running AUGUSTUS in *ab initio* mode resulted in 14,327 genes, whereas prediction with hints resulted in 17,277 genes.

ESTs and proteins from the Bergen *O. dioica* and our transcriptome assembly were used as evidence to run the MAKER pipeline (https://reslp.github.io/blog/My-MAKER-Pipeline/) that resulted in a set of 17,480 predicted genes. To finalize consensus gene structure, three predictions (weight = 1) were combined and subsequently refined with EvidenceModeler (EVM) v1.1.1 (Haas *et al.*, 2008) using PASA transcript assemblies (weight = 5) and proteins from the Bergen *O. dioica* (weight = 1) aligned to the genome with Exonerate v2.2.0, yielding 16,956 protein-coding genes. To predict UTRs and alternatively spliced isoforms, the EVM models were updated using two rounds of the PASA pipeline, which resulted in a final set of 18,485 transcript models distributed among 16,936 genes. Chromosomal coordinates were ported to our final assembly using the Liftoff tool (Shumate and Salzberg, 2020). The quality of the predicted gene models was assessed with BUSCO.

A draft annotation of the mitochondrial genome was obtained by submitting the corresponding scaffold (chr_Un12) as input to the MITOS2 mitochondrial genome annotation server (Bernt *et al.*, 2013; accessed May 28, 2020) with the ascidian mitochondrial translation table specified (Denoeud *et al.*,2010; Pichon *et al.*, 2019).

### Detection of coding RNAs

A translated alignment was used to detect known *O. dioica* genes available from GenBank using the TBLASTN software (Gertz *et al.*, 2006) with the options `-ungapped -comp_based_stats F` to prevent *O. dioica*’s small introns from being incorporated as alignment gaps, and `-max_intron_length 100000` to reflect the compactness of *dioica*’s genome. The best hits were converted to GFF3 format using BioPerl’s `bp_search2gff program` (Stajich *et al.*, 2002) before being uploaded to the ZENBU genome browser (Severin *et al.*, 2014). For some closely related pairs of genes that gave ambiguous results with that method, we searched for the protein sequence in our transcriptome assembly with TBLASTN, located the genomic region where the best transcript model hit was aligned, and selected the hit from the original TBLASTN search that matched this region. We summarized our results in Suppl. Table 6. For both searches, we used an *E*-value filter of 10^−40^. Genes marked as not found in the table might be present in the genome while failing to pass the filter.

### Detection of non-coding RNAs

To validate the results of cmscan on rRNAs, genomic regions were screened with a nucleotide BLAST search using the *O. dioica* isolate MT01413 18S ribosomal RNA gene, partial sequence (GenBank:KJ193766.1). 200-kbp windows surrounding the hits where then analysed with the RNAmmer 1.2 web service (Lagesen *et al.*, 2007). RNAmmer did not detect the 5.8S RNA, but we could confirm its presence by a nucleotide BLAST search using the AF158726.1 reference sequence. The loci containing the 5S rRNA (AJ628166) and the spliced leader RNA (AJ628166) were detected with the exonerate 2.4 software (Slater and Birney, 2005), with its affine:local model and a score threshold of 1000 using the region chr1:8487589-8879731 as a query.

### Whole-genome alignments

Pairs of genomes were mapped to each other with the LAST software (Kiełbasa *et al.*, 2011) version 1066. When indexing the reference genome, we replaced the original lowercase soft masks with ones for simple repeats (`lastdb -R01`) and we selected a scoring scheme for near-identical matches (`-uNEAR`). Substitution and gap frequencies were determined with `last-train` (Hamada *et al.*, 2017), with the alignment options - `E0.05 -C2` and forcing symmetry with the options `--revsym --matsym --gapsym`. An optimal set of pairwise one-to-one alignments was then calculated using `last-split` (Frith and Kawaguchi, 2015). For visualization of the results, we converted the alignments to GFF3 format and collated the colinear “match_part” alignment blocks in “match” regions using LAST’s command `maf-convert -J 200000`. We then collated syntenic region blocks (sequence ontology term SO:0005858) that map to the same sequence landmark (chromosome, scaffold, contigs) on the query genome with a distance of less than 500,000 bp with the custom script `syntenic_regions.sh` (supplementary Git repository). In contrast to the “match” regions, the syntenic ones are not necessarily colinear and can overlap with each other. The GFF3 file was then uploaded to the ZENBU genome browser.

### Nanopore read realignments

Nanopore reads were realigned to the genome with the LAST software as in the whole-genome alignments above. FASTQ qualities were discarded with the option `‒Q0` of `lastal`. Optimal split alignments were calculated with `last-split`. Alignment blocks belonging to the same read were joined with `maf-convert -J 1e6` and the custom script `syntenic_regions_stranded.sh`. The resulting GFF3 files were loaded in the ZENBU genome browser to visualize the alignments near gap regions in order to check for reads spanning the gaps.

### Analysis of sequence properties across chromosome-scale scaffolds

Each chromosome-scale scaffold was separated into windows of 50 Kbp and evaluated for GC content, repeat content, sequencing depth, and the presence of DpnII restriction sites. For chr 1, chr 2, and the PAR, windows corresponding to long and short chromosome arms were separated based on their positioning relative to a central gap region (chr 1 short arm: 1-5,191,657 bp, chr 1 long arm: 5,192,156-14,533,022 bp; chr 2 short arm: 1-5,707,009, chr 2 long arm: 5,707,508-16,158,756 bp; PAR short arm: 1-6,029,625 bp, PAR long arm: 6,030,124-17,092,476). Since none of our assemblies or sequencing reads spanned both the PAR and either sex-specific chromosome, the X and Y chromosomes were excluded from this analysis. For each of GC content, sequencing depth, repeat content, gene count, and DpnII restriction sites, the significance of the differences between long and short arms was assessed with Welch’s two-sided T test as well as a nonparametric Mann-Whitney test implemented in R (Suppl. Table 3). The results of the two tests were largely in agreement, but groups were only indicated as significantly different if they both produced significance values below 0.05 (*p* < 0.05).

## Data access

Sequence data was deposited to the ENA database (study ID PRJEB40135). Genome assembly and annotation were deposited to the NCBI (Accession number pending) and to Zenodo (DOI 10.5281/zenodo.4023777).

Custom scripts used in this study are available in GitHub (https://github.com/oist/oikGenomePaper).

## Acknowledgements

We would like to thank the DNA Sequencing Section and the Scientific Computing and Data Analysis Section of the Research Support Division at OIST for their support, Danny Miller for advices on Nanopore sequencing, Dan Rokhsar, Gene Myers, Ferdinand Marlétaz and Konstantin Khalturin for critical comments, Takeshi Onuma and Hiroki Nishida for sharing the OSKA2016 genome sequence prior publication, and Simon Henriet for sharing a DNA extraction protocol. MJM received funding as an International Research Fellow of the Japan Society for the Promotion of Science. This work was supported by core funding from OIST.

## Author contributions

Conceptualization: AB, CP, NML; data curation: AB, MJM, CP; formal analysis: AB, MJM, CW, TR, HCC, SK, JP, CP; investigation: AB, AM, MJM, YKT, AWL, CP; methodology: AB, CP; project administration: AB, CP; software: MJM, CW; supervision: CP, NML; validation: AB; visualization: AB, AM, MJM, YKT; writing – original draft: AB, AM, MJM, YKT, CP; writing – review & editing: AB, MJM, CP, NML.

## List of abbreviations

chr 1: autosomal chromosome 1
chr 2: autosomal chromosome 2
Kbp: Kilobase pairs
LGs: linkage groups
Mbp: Megabase pairs
PAR: pseudo-autosomal regions
XSR: X-specific region
YSR: Y-specific region

## Conflict of interests

The authors declare that they have no competing interests.

## Supplementary materials

**Supplementary Table 1:** Per-scaffold statistics

**Supplementary Table 2:** Contamination table

**Supplementary Table 3:** Statistics results for the analysis of sequence properties across chromosome-scale scaffolds

**Supplementary Table 4:** BUSCO scores

**Supplementary Table 5:** List of missing BUSCO genes in OKI2018_I69, OdB3 and OSKA2016 genome assemblies

**Supplementary Table 6:** Gene list uploaded to ZENBU

## Notes

https://github.com/oist/oikGenomePaper

https://doi.org/10.5281/zenodo.4023777

https://www.ebi.ac.uk/ena/browser/view/PRJEB40135

### Competing Interest Statement

The authors have declared no competing interest.

